# Nitazoxanide ameliorates OVA-induced airway inflammation in asthmatic mice via p38-MAPK/NFkB and AMPK/STAT3 pathways

**DOI:** 10.1101/2024.06.11.597851

**Authors:** Zhuyun Huang, Yanwen Zhang, Yudi Sun, Qian Wang, Zhonglan Men, Fuao Chu, Shuangyong Sun

## Abstract

Nitazoxanide has an anti-inflammatory effect, we clarified the ameliorative effect of nitazoxanide on asthmatic airway inflammation by conducting in vitro and in vivo experiments. In vitro, we assessed the effect of nitazoxanide on cytokine production by lipopolysaccharide-stimulated RAW 264.7 cells, as well as the diastolic effect of nitazoxanide on isolated rat airways. Nitazoxanide was found to have a diastolic effect on isolated tracheal spasms caused by spasmogenic substances, and to inhibit IL-6 and IL-1β production by RAW 264.7 cells. Meanwhile, nitazoxanide can inhibit the proliferation and migration of human bronchial smooth muscle cells (HBSMCs). In vivo, an ovalbumin (OVA)-induced asthma model was established in mice, and the airway resistance was measured by Whole Body Plethysmography (WBP) after inhalation of acetylcholine in mice, and the levels of IL-4, IL-6, IL-12, and IL-17 were detected in bronchoalveolar lavage fluid (BALF) of mice by ELISA and the inflammatory cells were counted. H&E staining was used to observe the changes in lung histopathology, and the expression of NFκB, MAPK, AMPK, and STAT3 in lung tissues was quantified using Western-blot. Nitazoxanide reduced inflammatory cell infiltration and goblet cell proliferation in the lungs of asthmatic mice. Moreover, the expression of IL-4, IL-5, and IL-6 in BALF was down-regulated in asthmatic mice. In addition, nitazoxanide could inhibit the expression of NFκB, MAPK, STAT 3 proteins and ascend the expression of AMPK in lung tissues. In conclusion, nitazoxanide could diastole airway smooth muscle and ameliorate OVA-induced airway inflammation in asthmatic mice via NFκB/MAPK and AMPK/STAT3 pathways.

## Introduction

Asthma is a highly heterogeneous disease[1] , characterized primarily by airway narrowing and airway inflammation. Asthma is mainly categorized into type 2 asthma and non-type 2 asthma[2] . Type 2 asthma is mainly due to an immune response triggered by exposure to external allergens including pollen, dust mites, pet hair, etc. , which leads to an upregulation of cytokine expression, such as IL-4 and IL-5, and ultimately leads to constriction of airway smooth muscle with eosinophilic airway inflammation[3]. Currently, the main therapeutic agents are glucocorticoids. Instead, type 2 asthma is triggered by air pollutants, viruses, bacteria, and obesity. The pathogenesis of non-type 2 asthma is complex and includes neutrophilic inflammation, type 1 and 3 immune responses, systemic inflammatory responses and metabolic abnormalities[4]. Glucocorticosteroids are poorly tolerated in patients with non-type 2 asthma, and there is a lack of effective therapeutic options[5] . At present, bronchodilators and anti-inflammatory drugs are the mainstay of asthma treatment[6] , whereas studies have shown that some asthmatics are desensitized to bronchodilators, such as beta agonists[7] . Meanwhile glucocorticoids have limitations in the treatment of non-type 2 asthma, so there is an urgent need for a new drug to solve the current dilemma[4] .

AMPK is an important metabolic perceptron that can influence cellular physiological processes by regulating intracellular energy levels[8] . In asthma treatment, activation of AMPK inhibits lung inflammation and also promotes cellular repair and resistance to oxidative stress, thereby alleviating asthma conditions[9]. STAT3 is an important transcription factor that can be activated by a variety of cytokines, and in asthma, activation of STAT3 can lead to an enhanced intracellular inflammatory response, thereby exacerbating the disease[10] . Studies have shown that activation of AMPK inhibits STAT3 activation and its downstream signaling, which influences airway smooth muscle proliferation[11] .The NFkB and sses such as inflammation, cell MAPK pathways are involved in asthma by regulating various cellular proceproliferation and cell survival[12,13] . Activation of these pathways by various stimuli, including allergens and pollutants, leads to activation of immune and inflammatory responses. Activation of the NFκB and MAPK pathways results in the production of proinflammatory cytokines and chemokines, leading to airway inflammation and constriction, which are characteristic symptoms of asthma[12,13] . Both pathways are also associated with airway remodeling, which is a permanent change in airway structure and is an essential cause of chronicity in asthma[14] .

TMEM16A (ANO1) is a Ca^2+^ activated Cl^-^ channel[15] , while studies have shown that TMEM16A in airway smooth muscle cells can regulate intracellular ion concentrations, thus affecting airway hyperresponsiveness[16] . Nitazoxanide, an FDA-approved anthelmintic, is also a TMEM16A antagonist, while studies have found potential anti-inflammatory activity[17] . In this study, we used both in vitro and in vivo experiments to observe the effect of nitazoxanide in the treatment of asthma and to explore its mechanism of action.

## Materials and methods

### Chemicals and reagents

Nitazoxanide was purchased from Shanghai yuanye Bio-Technology Co., Ltd (ShangHai, China), RAW264.7 cells, human bronchial smooth muscle cells (HBSMCs) and the corresponding medium were purchased from Procell Life Science&Technology Co. Ltd. (Wuhan, China) and were identified using immunofluorescence. Isoflurane was purchased from RWD Life science Co.,LTD (Shenzhen, China).Krebs-Henseleit nutrient solution (Procell, Wuhan, China),Immobilon-E PVDF membranes (0.45µm), OVA (grade V), Al(OH)_3_,DMSO, atropine sulfate, LPS (*Escherichia coli*) and acetylcholine were purchased from Sigma-Aldrich (St Louis, USA).PBS, RIPA lysate, 10% SDS-PAGE precast gel, Tris-glycine electrophoresis solution, Western blotting transmembrane solution, SDS Loading Buffer, Western blotting rapid containment solution and H&E staining solution were purchased from Beyotime (ShangHai, China). ELISA kits (IL-1β, IL-4, IL-5, IL-6) were purchased from Abcam (Cambridge, UK). BCA protein quantification kits were purchased from Thermo fisher scientific (Waltham, USA). AMPKα (F6), p-AMPKα (Thr172), STAT3 (124H6), p-STAT3 (Tyr750), p65-NFκB (L8F6), p-p65-NFκB ( Ser536), p38-MAPK (D13E1), p-p38-MAPK (28B10), β-Actin (8H10D10), goat anti-mouse secondary antibody, rabbit anti-mouse secondary antibody were purchased from Cell Signaling Technology (Boston, USA).

### Animals

Male SD rats and female BALB/mice used in this experiment were provided by SPF (Beijing) Biotechnology Co. All animals were housed in clean, pathogen-free animal houses with unrestricted access to standard laboratory food and water, on a 12-hour light/dark cycle, and regulated humidity (60%-80%) and temperature (22 ± 1 °C). The Animal Treatment and Use Committee of Tianjin Medical University School of Pharmacy accepted the animal experiments, which were conducted in strict accordance with ethical standards for institutional animal husbandry.

### RAW264.7 Cellular Anti-Inflammatory Assay

Nitazoxanide dissolved in DMSO. RAW264.7 cells were inoculated into 96-well plates according to 2×105 cells/well, 100 μL of cell suspension was added to each well, 50 μL of the corresponding concentration of the subject drug was added to the nitazoxanide group, and the corresponding volume of medium was added to the blank and model groups. After 1h of incubation, 50μL of LPS (final concentration of LPS was 100 ng/mL) was added, and the control group was added with the corresponding volume of medium. After 24h of incubation in the incubator, the supernatant was centrifuged at 1000rpm for 5min, and the supernatant was taken to detect the levels of IL-1β and IL-6 (Abcam, USA) and calculate the IC50.

### Proliferation and migration of HBSMCs

HBSMCs and appropriate complete medium were purchased from Procell Life Science&Technology Co., Ltd. (Wuhan, China) and identified using immunofluorescence method. The proliferation experiments were divided into blank group and nitazoxanide group (10 and 25 μM), after the HBSMCs were passaged to the third generation, 300 cells/well were inoculated into 96-well plates, and 100 μL of cell suspension was added to each well, 100 μL of the corresponding concentration of the test drug was added to the nitazoxanide group, and the corresponding volume of medium was added to the blank group, and 20 μL of MTS reagent was added to each well on days 1, 3, and 6, and incubated for 1 h at 37°C, and measure the absorbance of each well at 490nm by using an enzyme meter.

In cell migration experiments, cells were inoculated in 6-well plates at 2 × 10^6^ cells/well, and after cell attachment, the medium was discarded after vertical scoring by using a 200 μL lance tip, washed with PBS, and medium containing nitazoxanide (10, 25 μM) was added, the blank control group was added with the corresponding volume of medium. The degree of cell migration was quantified by measuring the area of the cell-free zone using Image J software and a fluorescence inverted microscope (OLYMPUS, Inc., Tokyo, Japan).

### Strips of trachea

Rats were executed by using a small animal respiratory anesthesia machine (RWD Life science Co., LTD, Shenzhen, China) with isoflurane overdose anesthesia, and the trachea was removed and placed into a flat dish containing Krebs-Henseleit nutrient solution, after the connective tissues around the trachea were cut, the trachea was cut into a spiral strip with a length of 4 mm and connected to a tension transducer ( Chengdu Taimeng Software Co., Ltd, Chengdu, China), and then the fixed tracheal strips were immobilized by placing them into an ex vivo organ bath (Chengdu Taimeng Software Co., Ltd, Chengdu, China) containing 20 mL of K-H solution, a mixture of 5% carbon dioxide and 95% oxygen was continuously introduced. A constant temperature of 37°C was maintained, the load was 1.5 g, and the Krebs-Henseleit solution was changed 2-3 times per hour, the specimens were stabilized in the fixed bath for about 1 h and then traced. After the isolated tracheal strips were stabilized, 10 μM acetylcholine (Ach) was added to the bath, causing the tracheal strips to contract, after the contraction was stabilized, a gradient concentration of nitazoxanide was added at 3 min intervals (nitazoxanide was dissolved in DMSO) to a final concentration of 0.005-50 μM, the corresponding volume of DMSO was added simultaneously to the ACh group. 1 μM atropine sulfate was added to completely diastole the tracheal strips after the trachea was stabilized in both groups.

### OVA-induced asthma in mice

#### Programmatic and pharmacological treatments

Thirty female BALB/c mice were acclimatized and fed for one week, and then divided into two batches, the first batch was randomly taken out of 10 as a blank control group, and the remaining 20 mice were used to prepare the asthma model, respectively, the model group, the nitazoxanide group (dosage: 10 mg/kg), and 10 animals in each group. The model mice were sensitized by intraperitoneal injection of 0.2 mL of sensitizing solution, 1 mL of phosphate buffer solution (PBS) including 0.25 mg of OVA and 5 mg of Al (OH)_3_ per milliliter of sensitizing solution on days 0, 7, 14, respectively, while the normal group was injected with an equal volume of PBS.

During days 21 to 23 of sensitization, mice in the model and nitazoxanide groups were placed in a homemade nebulizer box, and then 5% OVA was nebulized by using a nebulizer (PARI, Starnberg, Germany) for 30 minutes for nebulized inhalation, and mice in the normal group were nebulized for the same time with PBS. Mice were administered 0.2 mL of the solvent by gavage 30 minutes before the nebulization excitation each day, in which only the corresponding volume of solvent was given by gavage in the normal and model groups once a day for 3 consecutive days.

#### Measurement of airway hyperresponsiveness

Airway responsiveness was measured 24 hours after the last excitation in each group of mice. Mice were placed in a WBP (EMMS, London, UK), and after measurement of the basal value of the expiratory interval (Penh), they received excitation with nebulized acetylmethacholine, which was administered at concentrations of 0, 6.25, and 12.5 mg/mL, with a nebulized dose of 150 μL per session for 60 s. After completion of nebulization at each concentration, 3 min Penh values were recorded to compare airway responsiveness in each group of mice.

#### Tissue sampling and evaluation of BALF

Once the airway hyperresponsiveness test in mice was completed, the mice were anesthetized using an anesthesia machine, and blood was taken from the orbits of the mice, then over-anesthetized and executed; he chest cavity was opened, the left lung was ligated, and the right lung was lavaged with 0.5 mL of pre-cooled PBS three times in the lungs of the mice. Then the BALF was centrifuged at 3000 rpm for 5 min, and after centrifugation, the supernatant was dispensed and stored at -20 °C; the precipitated cells were resuspended with 1.5 mL of pre-cooled PBS. The cells detected including leukocytes and lymphocytes were counted using a cell analyzer (Mindray, Shenzhen, China). Levels of IL-4, IL-5, and IL-6 in BALF supernatants were determined using ELISA kits.

#### Western blotting

Take 50mg of frozen lung tissue, add 500µL of RIPA lysate, ultrasonically broken and centrifuged to get the supernatant, use BCA protein quantification kit for protein quantification, add Loading Buffer proportionally and boil for 10min, separate by 10% SDS-PAGE, transfer the protein onto a PVDF membrane (0.45µm), and close it using rapid Proteins were transferred onto PVDF membrane (0.45µm) and sealed with rapid sealing solution for 10min, primary antibody was incubated at 4°C overnight, secondary antibody was incubated at room temperature for 1 hour, chemiluminescence was used to develop the color, and the proteins were imaged on a gel imager (Blolight, Guangzhou, China), the protein bands were analyzed by Image J.

#### Histopathological examination

The samples of mouse lung tissue were taken and fixed in 4% formaldehyde, embedded in paraffin, and cut into 4 μm sections. Sections were stained using hematoxylin and eosin (H&E). A Leica laser microdissection system was used to capture the images (Leica Biosystems, Heidelberg, Germany). Image Scope was used to capture images of the sections for analysis (OLYMPUS, Inc., Tokyo, Japan).

### Statistical analysis

Statistical analysis was performed with SPSS software, and the measurement information was expressed as mean ± standard deviation, and One-Way ANOVA was used for comparison between groups, LSD test was used for chi-square, and Dunnett-t test was used for non-chi-square, and the difference of P<0.05 was statistically significant.

## Results

### Nitazoxanide inhibited LPS-induced expression of IL-1β, IL-6 in RAW264.7 cells

One of the main symptoms of asthma is bronchial inflammation. We first evaluated the in vitro anti-inflammatory effect of nitazoxanide. In the LPS-induced inflammation model of RAW264.7 cells, nitazoxanide was able to significantly down-regulate the expression of two inflammatory factors, IL-1β and IL-6. The calculated IC50 of nitazoxanide on IL-1β and IL-6 were 10.08 μM and 16.9 μM, respectively (Fig 1A, B).

**Figure 1.**
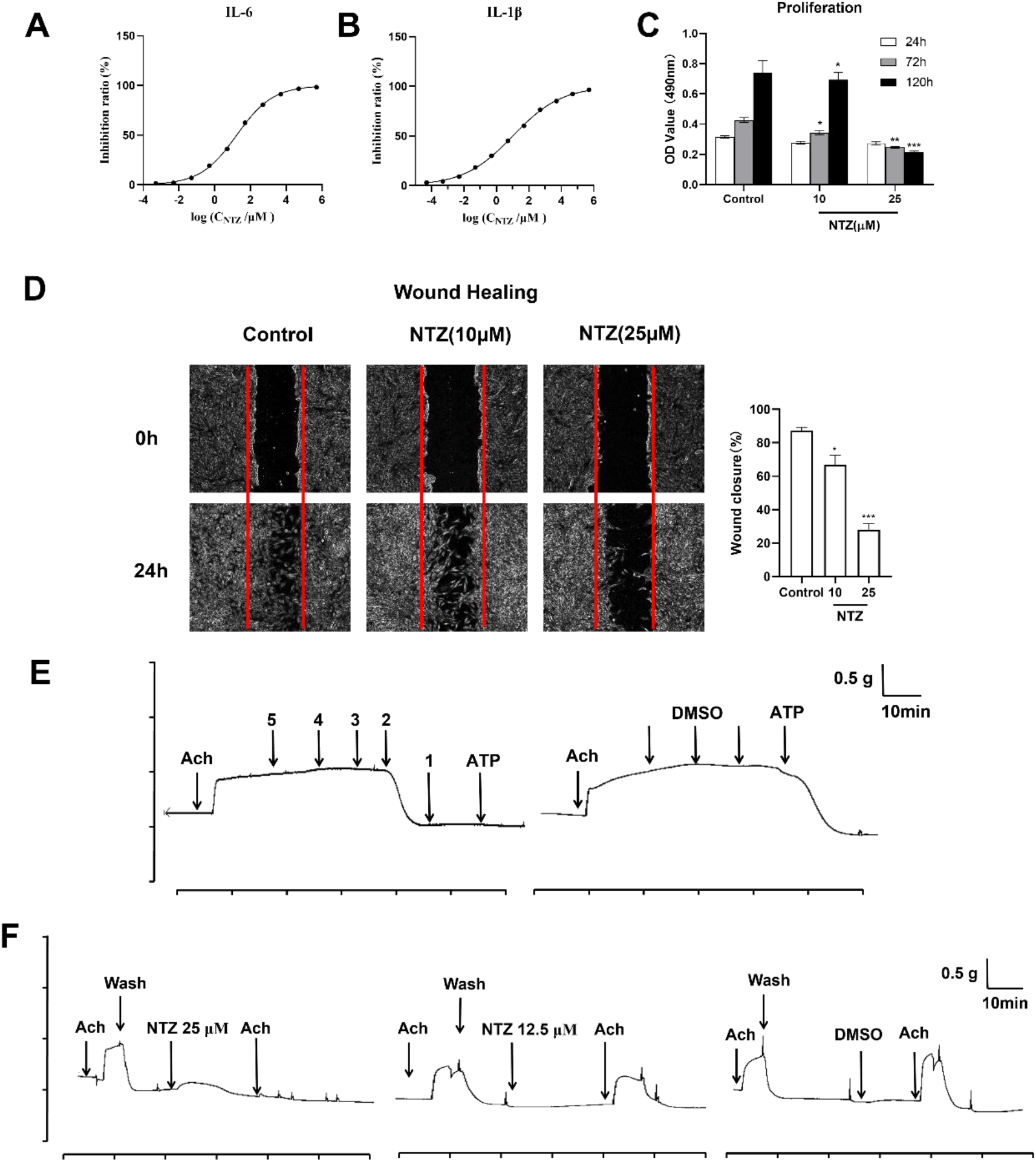
In vitro pharmacodynamic assay of nitazoxanide. A and B, Quantitative efficacy curves of nitazoxanide on IL-1β, IL-6 in LPS-induced RAW264.7 cells. C, Proliferation of HBSMCs were inhibited after being treated with nitazoxanide for 24, 72, and 120h, * P<0.05, ** P<0.01, *** P<0.001 vs control. D, Nitazoxanide inhibits migration of HBSMCs, raw images on the right, pooled data on the left, * P<0.05, *** P<0.001 vs control. E, Nitazoxanide dose-dependently diastole Ach-induced in isolated rat trachea. 5, 0.005 μM, 4, 0.05 μM, 3, 0.5 μM, 2, 5 μM, 1, 50 μM, ATP for atropine. F, Pretreatment of nitazoxanide inhibited Ach-induced contraction of tracheal strips with Ach.

### Nitazoxanide inhibits the proliferation and migration of HBSMCs

The proliferation and migration of HBSMCs are closely related to airway remodeling, and we further evaluated the proliferation and migration of HBSMCs under different treatment durations and different administration concentrations of nitazoxanide. The results showed that both 10 μM and 25 μM nitazoxanide effectively inhibited the proliferation and migration of HBSMCs in a dose-dependent manner (Fig 1C, D).

### Nitazoxanide dose-dependently relaxes Ach-treated tracheal strips and pretreatment with nitazoxanide inhibits contraction of Ach-treated tracheal strips

The contraction of bronchial smooth muscle causes tracheal stenosis and dyspnea. In order to assess the effect of nitazoxanide on spasmogenic substance-induced tracheal constriction, we used isolated tracheal strips to confirm the effect of nitazoxanide on tracheal constriction, and the results showed that nitazoxanide was able to dose-dependently relax Ach-treated tracheal strips (Fig 1E).

Acute asthma attacks are usually characterized by severe bronchoconstriction, which leads to respiratory distress. To verify whether nitazoxanide prevents airway constriction caused by spasmogenic substances, we measured the effect of Ach (10 μM) on airway strip constriction in the presence of nitazoxanide pretreatment. The results showed that Ach on tracheal bar constriction was relieved after nitazoxanide pretreatment (Fig 1F).

### Nitazoxanide alleviates airway hyperresponsiveness as well as lung inflammation in OVA-induced asthmatic mice

After the mice were sensitized with OVA+Al (OH)_3_ for 3 weeks and then stimulated with 5% OVA for 3 consecutive days and then with acetylmethacholine at 6.25 mg/mL versus 12.5 mg/mL, Penh values were significantly higher compared with the control group. Penh values were significantly lower in the nitazoxanide group compared to the model group (Fig 2A).

**Figure 2.**
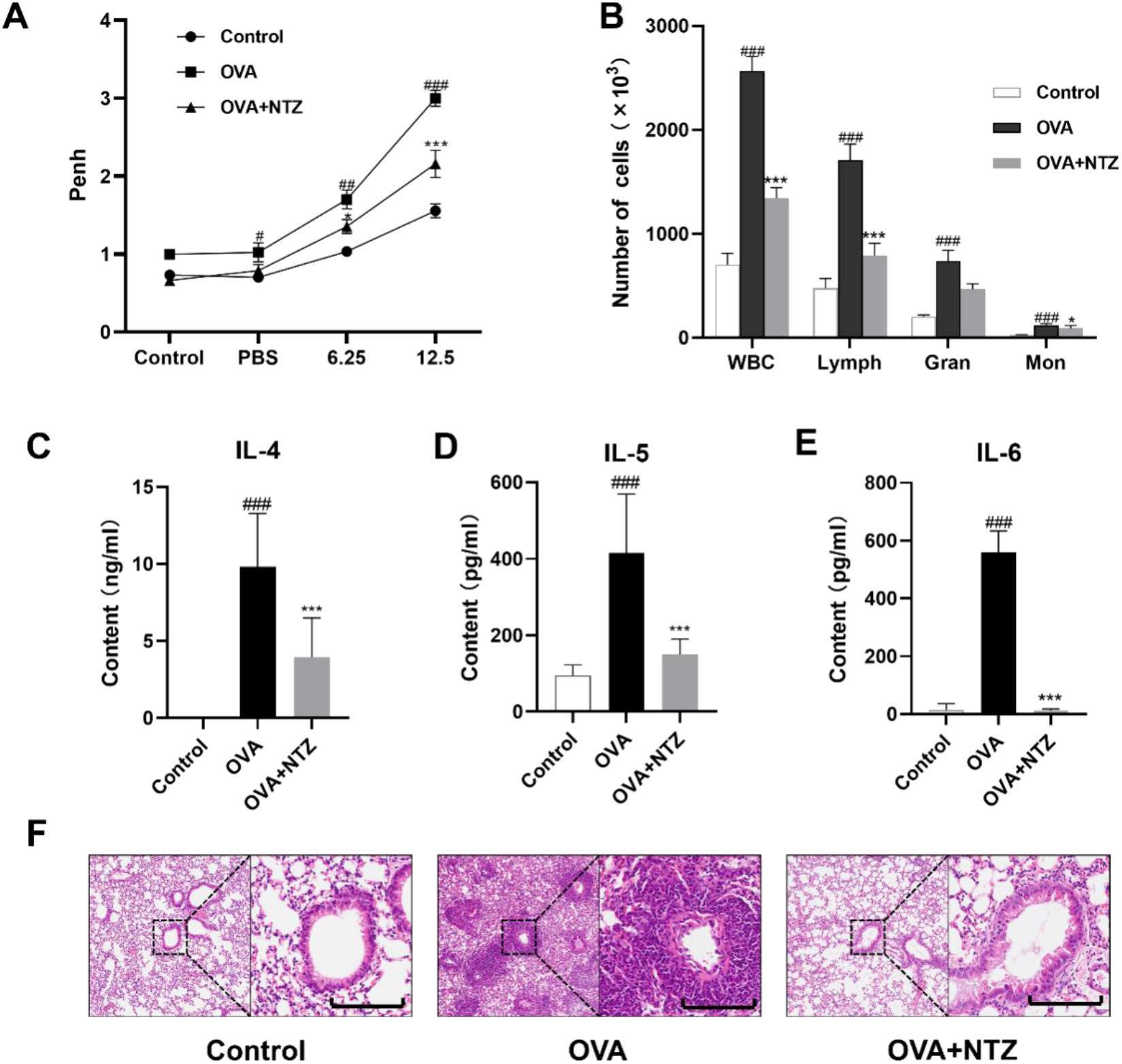
Therapeutic effect of nitazoxanide on OVA-induced asthma in mice. A, Magnitude of Penh values in each group after mice were stimulated with acetylmethacholine at 6.25 mg/mL versus 12.5 mg/mL.B, Nitazoxanide reduced the inflammatory cellular content of the BALF, C, D, E, Nitazoxanide intervention decreased IL-4, IL-5, and IL-6 levels in BALF. f, H&E staining of mouse lungs in each group, scale bar, 100 μm.* P<0.05, *** P<0.001 vs model, # P<0.05, ## P<0.01, ### P<0.001 vs control.

To determine the severity of inflammation in the lungs of asthmatic mice, we measured the contents of BALF in each group of mice. First, we counted inflammatory cells in the BALF, which showed that inflammatory cells were significantly elevated in the lungs of mice that had undergone OVA induction compared with the control group, whereas mice treated with nitazoxanide had reduced levels of inflammatory cells in the lungs (Fig 2B). The supernatant of BALF was then used to measure the content of cytokines IL-4, IL-5, IL-6, which were similar to the levels of inflammatory cells, and after nitazoxanide treatment, the levels of cytokines IL-4, IL-5, and IL-6 were reduced (Fig 2C, D, and E).

H&E staining of the lungs of OVA-induced mice showed significant pathologic structural changes in the lungs, with a large infiltration of inflammatory cells as well as hyperplasia of goblet cells, which were significantly alleviated by treatment with nitazoxanide (Fig 2F).

### Nitazoxanide upregulates AMPK expression and inhibits NFkB, p38-MAPK and STAT3 protein expression in lung tissues

It has been shown that AMPK, NFκB, p38-MAPK and STAT3 proteins are all associated with airway remodeling as well as airway inflammation in asthma. We investigated the effect of nitazoxanide on the expression levels of these proteins. Nitazoxanide treatment increased the expression of AMPK and decreased the expression of NFκB, p38-MAPK, and STAT3 proteins in the lung tissues of mice. The results suggest that nitazoxanide reduces the level of lung inflammation and airway hyperresponsiveness by activating the AMPK pathway and inhibiting the NFκB, p38-MAPK and STAT3 pathways (Fig 3).

**Figure 3.**
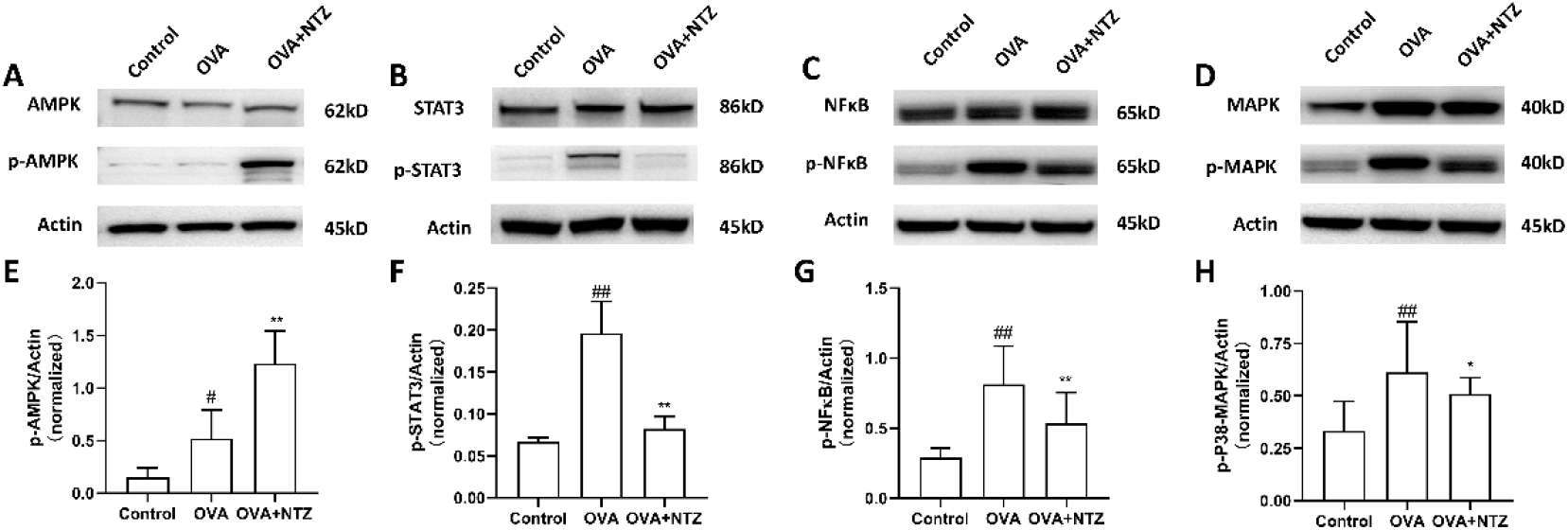
Nitazoxanide up-regulated the expression of AMPK and inhibited the expression of NFκB, p38-MAPK and STAT3 proteins in lung tissue. * P<0.05, ** P<0.01 vs model, # P<0.05, ## P<0.01 vs control.

## Discussion

Asthma is a chronic airway disease characterized primarily by airway hyperresponsiveness, airway inflammation, and airway remodeling[18] . Th2 cytokines play a key role in airway inflammation, and during an asthma attack, the excessive release of IL-4 and IL-6 from Th0 cells causes Th2 and Th17 to produce a number of pro-inflammatory factors, which promote the activation and proliferation of inflammatory cells eosinophils, macrophages, and mast cells[3] .

Nitazoxanide is a thiazolylated compound that has long been used as an antiparasitic against protozoa, nematodes, and trematodes[19]. Nitazoxanide increases the phosphorylation of PKR and induces eIF2a, which ultimately inhibits the translation of viral RNA, resulting in therapeutic effects against hepatitis B and hepatitis C[20]. Nitazoxanide is also able to target the Wnt/β-catenin signaling pathway for the treatment of cancer[21]. Recently, nitazoxanide has been found to have mitochondrial uncoupling effects, and mitochondrial uncoupling agents have been shown to inhibit smooth muscle cell proliferation and migration[22]. Meanwhile, nitazoxanide, a TMEM16A inhibitor, can regulate mucin secretion and diastolic airway smooth muscle[16].

Due to the inhibitory effect of nitazoxanide on TMEM16A, we further verified the diastolic effect of nitazoxanide on airway smooth muscle through experiments, and discovered that nitazoxanide had a preventive effect on bronchospasm. Meanwhile, we confirmed that nitazoxanide could inhibit the proliferation and migration of bronchial smooth muscle cells through the proliferation and migration experiments of HBSMCs. In conclusion, nitazoxanide can affect airway remodeling. We further established the LPS-induced inflammation model of RAW264.7 cells and evaluated the anti-inflammatory effect of nitazoxanide, and the results showed that nitazoxanide could improve the inflammation of LPS-induced RAW264.7 cells, and significantly down-regulated the expression of IL-1β and IL-6, which indicated that nitazoxanide had certain anti-inflammatory effect.

Based on the above in vitro experiments, we further conducted in vitro experiments using an OVA-induced asthma mouse model. After the intervention of nitazoxanide, the airway hyperresponsiveness of asthmatic mice was reduced, which was consistent with the results of our in vitro experiments. After evaluating the severity of lung inflammation, the results showed that nitazoxanide was able to ameliorate lung inflammation in OVA-induced asthmatic mice. These indicate that nitazoxanide possesses both bronchodilator and anti-inflammatory effects in the anti-asthmatic process. Acute asthma attacks require prompt use of bronchodilators to dilate the airways, while chronic asthma requires anti-inflammatory drugs for long-term treatment, so nitazoxanide has the potential to be used as a therapeutic agent for either acute asthma or chronic asthma. However, it has been found that the water solubility of nitazoxanide is poor, and oral administration needs to be made into eutectic to have high bioavailability[23]. Most of the asthma treatments are administered locally through the respiratory system, so there is a need to improve the water solubility of nitazoxanide for respiratory administration, thus reducing the amount delivered.

Recent studies have shown that nitazoxanide inhibits STAT3 protein expression^[24]^ . In mice, we found that nitazoxanide down-regulated the expression of STAT3, p38-MAPK and NFκB proteins, and up-regulated the expression of AMPK proteins. STAT3, p38-MAPK, NFκB, and AMPK signaling are critical in the pathology of asthma. Activation of STAT3, p38-MAPK and NFκB leads to proliferation and migration of vascular and bronchial smooth muscle cells[8,25,26]. Activation of AMPK can dilate bronchioles and inhibit airway smooth muscle proliferation[27] . In the present study, we found that nitazoxanide activated AMPK in lung tissues, and we hypothesized that nitazoxanide could dilate bronchioles by activating AMPK, thereby reducing airway hyperresponsiveness. Up-regulation of AMPK can lead to down-regulation of STAT3 expression[28] and downregulation of STAT3 inhibits IgE-mediated proliferation of human airway smooth muscle cells[29]. In addition, reduction of STAT3 can inhibit IgE-mediated proliferation of human airway smooth muscle cells. Furthermore, downregulation of STAT3 can also reduce the infiltration of immune cells in the airway[30] Cao et al. reported that inhibition of p38-MAPK reduced STAT3 activation and suggested that STAT3 may be a substrate for p38-MAPK [31]. p38-MAPK is a member of the MAPK family involved in a variety of asthma cellular activities, such as cuprophagocytosis, smooth muscle cell migration and cytoskeletal remodeling, and epithelial-mesenchymal transition (EMT) [31–33]. Therefore, nitazoxanide can also inhibit airway remodeling by inhibiting p38-MAPK.

NFκB is a nuclear transcription factor involved in the regulation and expression of genes associated with inflammatory factors[34] . In allergic asthma, activation of NFκB can lead to the transition of T cells to Th2 cells and subsequent production of Th2 cytokines (IL-4 and IL-5)[35]. At the same time, upregulation of NFκB can also activate gene expression of IL-6[36] . It has been reported that IL-6 can be regulated by p38-MAPK and STAT3, and inhibition of IL-6 can also downregulate the expression of p38-MAPK and STAT3[34,37] . From our results, nitazoxanide is able to diminish the expression of IL-5, IL-4 and IL-6 in lung tissues, as well as the expression of NFκB.

In summary, nitazoxanide inhibits the expression of inflammatory factors and the aggregation of inflammatory cells in the lungs by mediating the AMPK, STAT3, p38-MAPK and NFκB signaling pathways, which ultimately alleviates inflammation and airway remodeling. Meanwhile, nitazoxanide exerted a diastolic effect on airway smooth muscle through the inhibition of TMEM16A. We hypothesize that nitazoxanide could be a candidate for the treatment of asthma or could be used as a lead compound to develop anti-asthma drugs with better water solubility and effect.

## Acknowledgments

The authors are grateful to Tianjin Medical University for the provision of experimental sites

## CRediT authorship contribution statement

Zhuyun Huang: Writing – original draft. Yanwen Zhang: Writing – review & editing. Yudi Sun: Writing – review & editing. Qian Wang: Writing – review & editing. Zhonglan Men: Writing – review & editing. Fuao Chu: Writing – review & editing.

